# Elexacaftor/Tezacaftor/Ivacaftor alters branching morphogenesis of the mouse embryonic lung

**DOI:** 10.1101/2021.11.01.466814

**Authors:** Mickaël Lhuillier, Laura Aoust, Elise Dreano, Marie-Laure Franco-Montoya, Kim Landry-Truchon, Nicolas Houde, Stéphanie Chhun, Alexandre Hinzpeter, Aleksander Edelman, Christophe Delacourt, Lucie Jeannotte, Isabelle Sermet-Gaudelus, Alice Hadchouel

## Abstract

**Introduction:** CFTR modulators triple combo-therapy Elexacaftor/Tezacaftor/Ivacaftor (ETI) has proven to clinically benefit homozygous and heterozygous F508del patients. As a result, an increasing number of pregnancies is expected. Studies of the potential impact of these modulators on the development of the foetus are mandatory.

**Materials:** We used the early mouse embryonic lung organ culture model to analyse *ex vivo* the lung branching process and the relative expression of *Fgf10*, *Fgfr2IIIb*, *Shh*, and *Hhip* development regulator genes in different conditions: standard culture medium, treatment with ETI or with Forskolin ± Inh172. Development of lung branching and distal bud caliber were evaluated in lung explants from heterozygous F508del *Cftr*^tm1Eur/+^ and control *Cftr*^tm1Eur+/+^ (WT) mouse embryos at E12.5 during pseudo-glandular stage.

**Results:** Exposure to ETI of the *Cftr*^tm1Eur/+^ and WT lung explants induced a significant decrease in lung branching after 48h culture and the percentage of terminal bud dilations was significantly increased. These results were recapitulated by cAMP-dependent CFTR continuous activation by Forskolin and reversed by addition of Inh172.

ETI induced a significant decrease in *Fgf10*, *Fgfr2IIIb*, *Shh* and *Hhip* expression in lung explants of both E12.5 *Cftr*^tm1Eur/+^ and WT embryos treated with ETI for 72h.

**Conclusion:** Our results provide evidence that the triple association Elexacaftor/Tezacaftor/Ivacaftor alters lung branching morphogenesis of WT and heterozygous F508del mouse embryos during the pseudo-glandular stage. Those results argue for a close monitoring of pregnancies in patients treated with these drugs.

**Plain Language:** *Introduction:* The triple combo-therapy Elexacaftor/Tezacaftor/Ivacaftor (ETI) improves homozygous and heterozygous F508del patients. As a result, an increasing number of pregnancies is expected. Studies of this treatment on the development of the foetus are lacking. We incubated lungs of murine foetus not carrying CFTR mutation or F508del heterozygous. We show that ETI induces significant defect of lung development and the formation of cysts. These results are at least partly due to CFTR activation. Those results argue for a close monitoring of pregnancies in patients treated with these drugs.

## Introduction

Cystic Fibrosis (CF) is the most common monogenic autosomal recessive disease of people of Caucasian descent with an estimated incidence between 1/3000 to 1/6000 births in European populations [1, 2]. CF is caused by mutations in the *Cystic Fibrosis Transmembrane Conductance Regulator* (*CFTR*) gene, which leads to defective expression and/or function of the CFTR protein. The most frequent mutation in the Caucasian population is the Phe508del mutation (F508del thereafter). In the respiratory tract, CFTR dysfunction is associated with altered mucus clearance, chronic lung inflammation, recurrent respiratory infections, and ultimately respiratory failure [3].

CFTR modulators triple combo-therapy Elexacaftor/Tezacaftor/Ivacaftor (ETI hereafter) has proven to clinically benefit homozygous and heterozygous F508del patients [4–7] and to improve CF patient’s fertility. As a result, young women with CF are more likely to become pregnant and give birth [8]. With the lack of evidence of safety during pregnancy, studies aiming to understand the potential impact of these modulators on the development of the foetus are mandatory. Until now a limited number of pregnancies upon CFTR modulators, (mainly Ivacaftor or Lumacaftor-Ivacaftor) [9–14], and very few upon ETI, occurred and showed no alarming signals [15].

In the human developing lung, CFTR expression is detected from the 12^th^ week of gestation [16] and is involved in lung fluid secretion and airway budding [17]. This is suggested by observations in pigs, which showed that invalidation of CFTR alters pseudo-glandular development inducing a significant reduction of the caliber of the trachea and the proximal bronchi together with the formation of hypo-distended airway budding [18]. In animal reproductive models, Ivacaftor, Tezacaftor and Elexacaftor cross the placenta, and the mean foetal lung concentration ratio is very high, around 200% of the plasma, much higher than into brain or cerebrospinal fluid [13, 19].

We asked whether ETI might impact foetal lung organogenesis, and more specifically if it might affect lung branching and distal bud formation in mice carrying the +/F508del heterozygous genotype. To reach this aim, we investigated the effect of ETI on the development of embryonic lung during pseudo-glandular stage from heterozygous +/F508del *Cftr* mice and more specifically lung branching and distal buds’ formation. We used the early mouse embryonic lung organ culture model to analyse *ex vivo* the lung branching process and further dissect the molecular mechanisms involved.

## Material and methods

### Animal experiments

Experiments were performed according to the guidelines of European directive 2010/63/UE with the approbation of ethics committee for animal experiments of Paris Descartes University (CEEA n°34).

C57Bl6j male and female wild type (WT) mice were obtained from Charles River (Saint-Germain-Nuelles, France). Heterozygous male and female mice carrying the F508del mutation in the *Cftr* gene (*Cftr*^tm1Eur^ mouse) [20] were obtained from CNRS-UPS44 “Transgenesis and storage of animal models” at Orléans (France). Age of the embryos was estimated by considering the morning of the day of vaginal plug as the first half day of pregnancy (E0.5). Pregnant female mice were killed by cervical dislocation and lungs were collected from embryos after 12 days of gestation (E12.5) corresponding to the human pseudo-glandular stage (5 to 17 weeks of gestation). Tail tissue was collected for genotyping by PCR.

### Mouse embryo lung explant cultures

Lung explants were collected from E12.5 mouse embryos and cultured in serum-free Dulbecco’s Modified Eagle Medium (DMEM/F12 (Gibco 31331093, Thermofischer Scientific, Illkirch, France) for 72h on porous membranes (Whatman ® Nucleopore™ Track-Etched Membranes WHA150446, Sigma Aldrich, St. Quentin Fallavier, France) in 4-well plates [21]. Pictures of the explants were taken at baseline (T0) and after 24 hours (T24), 48 hours (T48) and 72 hours (T72) of culture. The number of terminal buds at the periphery of each sample was counted and the branching growth was estimated by the relative ratio (Branches T24, T48 or T72−Branches T0) × 100/Branches T0 as previous described by Boucherat *et al* [22]. It was compared to the DMEM/F12 control group, at each time point, in percentage of variation from this reference group.

The diameter of the terminal buds of the explants was measured after 72h of culture using the ImageJ software. A terminal bud dilation was defined by a diameter of the terminal bud greater than two standard deviations (SD) from the mean diameter of the DMEM/F12 control group. The results were reported as the ratio of the number of dilations in the treated group to that reported in the DMEM/F12 control group.

To modulate CFTR protein activity, culture medium was supplemented with CFTR modulators combination at the following concentrations:

- Elexacaftor (S8851, Selleckchem, Souffelweyersheim, France) diluted in DMSO (10μM final),
- Tezacaftor (S7059, Selleckchem, Souffelweyersheim, France) diluted in DMSO (10μM final),
- Ivacaftor (S1174, Selleckchem, Souffelweyersheim, France) diluted in DMSO (100nM final),
- Forskolin (F6886, St. Quentin Fallavier, France) diluted in EtOH 100% (5μM final) was used to increase cAMP intracellular content and activate CFTR phosphorylation,
- CFTR inhibitor Inh-172 (C2992, Sigma Aldrich, St. Quentin Fallavier, France) diluted in DMSO (5μM final) was used to specifically inhibit CFTR.

### Real-time quantitative PCR analysis

Expression of *Cftr* expression was monitored during mouse lung development by RT-qPCR assays during gestation at E12.5, E15.5 and E18.5 and after birth at P0, P7, P15 and P30 in WT specimens (**See Online supplement**).

RT-qPCR expression analyses of *Fgf10*, *Fgfr2IIIb*, *Shh*, and *Hhip* were performed with RNA from E12.5 lung explants snap-frozen after 72h of culture (**See Online supplement**).

### Statistical analyses

Data were expressed as mean and Standard Error of the Mean (s.e.m). They were analysed with repeated measures ANOVA, and two groups comparison were performed by Mann-Whitney U test using GraphPad Prism software (GraphPad Software INC, San Diego, CA). A significance level inferior to 5% (p<0.05) was considered statistically significant.

## Results

### The F508del mutation does not alter lung branching morphogenesis in mice

In WT mice, *Cftr* expression was detected throughout lung formation from E12.5 onwards with a maximum level at E18.5 just before birth (**Figure S1**).

There was no significant difference between lung explants from WT *Cftr*^+/+^, F508del heterozygous *Cftr*^tm1Eur/+^ or F508del homozygous *Cftr*^tm1Eur/tm1Eur^ embryos at E12.5 neither for lung branching nor for terminal buds’ dilations (**Figure 1**, **Table 1**). Lung branching increased similarly in the three groups at each time point, up to 302% ± 27% in the WT group, 263% ± 15% in the *Cftr*^tm1Eur/+^ group, and 281% ± 20% in the *Cftr*^tm1Eur/tm1Eur^ group at T72 (Not Significant (NS)). Moreover, there was no significant difference in the percentage of terminal buds’ dilations after 72h of culture between the three groups. Those results demonstrate that carrying one or two *Cftr* F508del variant alleles does not have major impact on lung branching morphogenesis.

**Figure 1.**
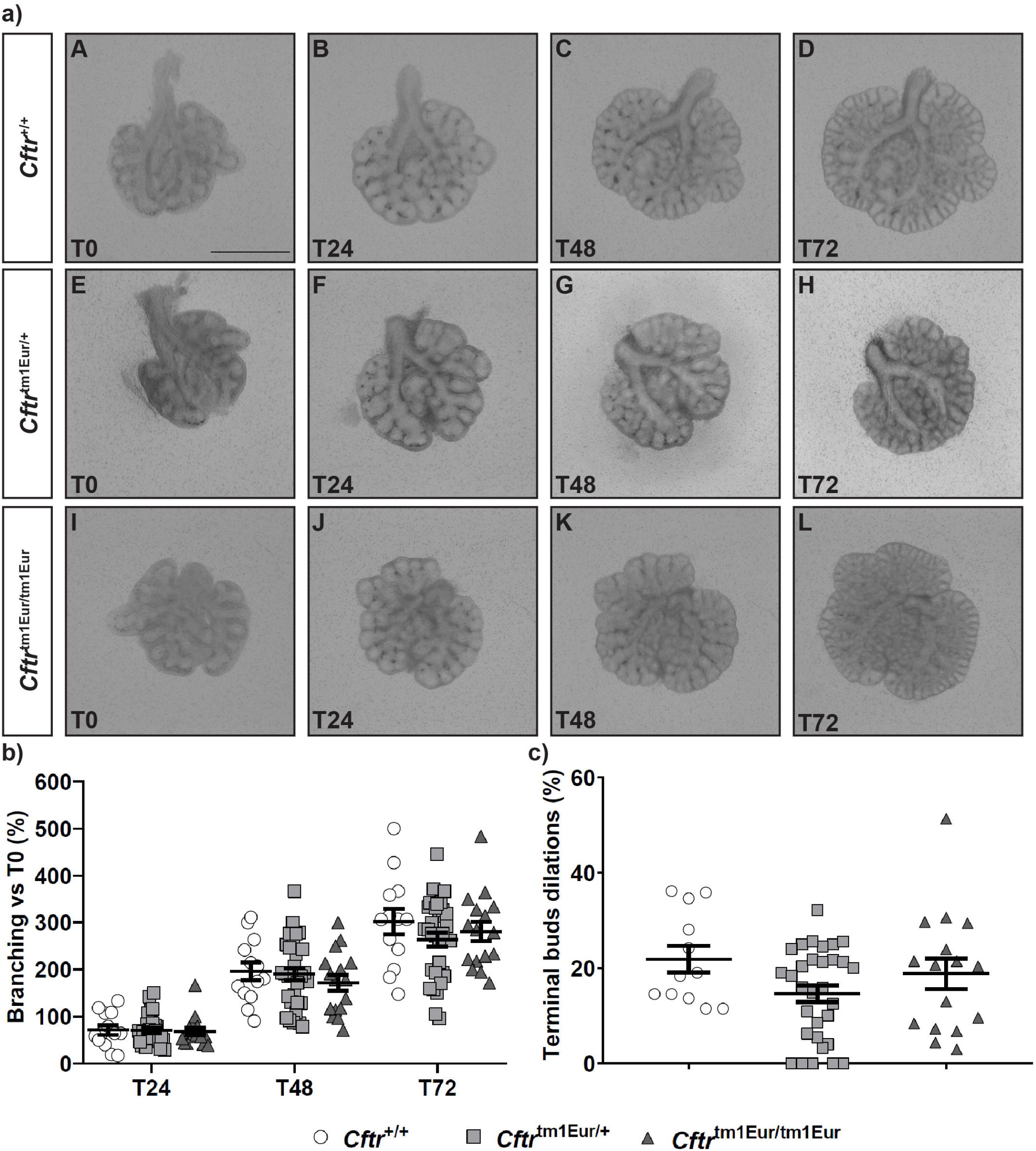
*Ex vivo* lung explants from E12.5 *Cftr*^tm1Eur^ mouse embryos. A. Lung explants from WT (A-D)(n=12), *Cftr*^tm1Eur/+^ (E-H)(n=31), and *Cftr*^tm1Eur/tm1Eur^ (I-L)(n=16) embryos cultured for 3 days in DMEM/F12. Scale bar: 1mm. B. Number of terminal buds at T24, T48 and T72, expressed as the percentage in branching increase compared to baseline (T0). C. Number of terminal buds’ dilations at T72, expressed as the percentage of the dilations increase compared to mean number of terminal buds’ dilations in the media group. Data are expressed as mean ± s.e.m.

**Table 1.**
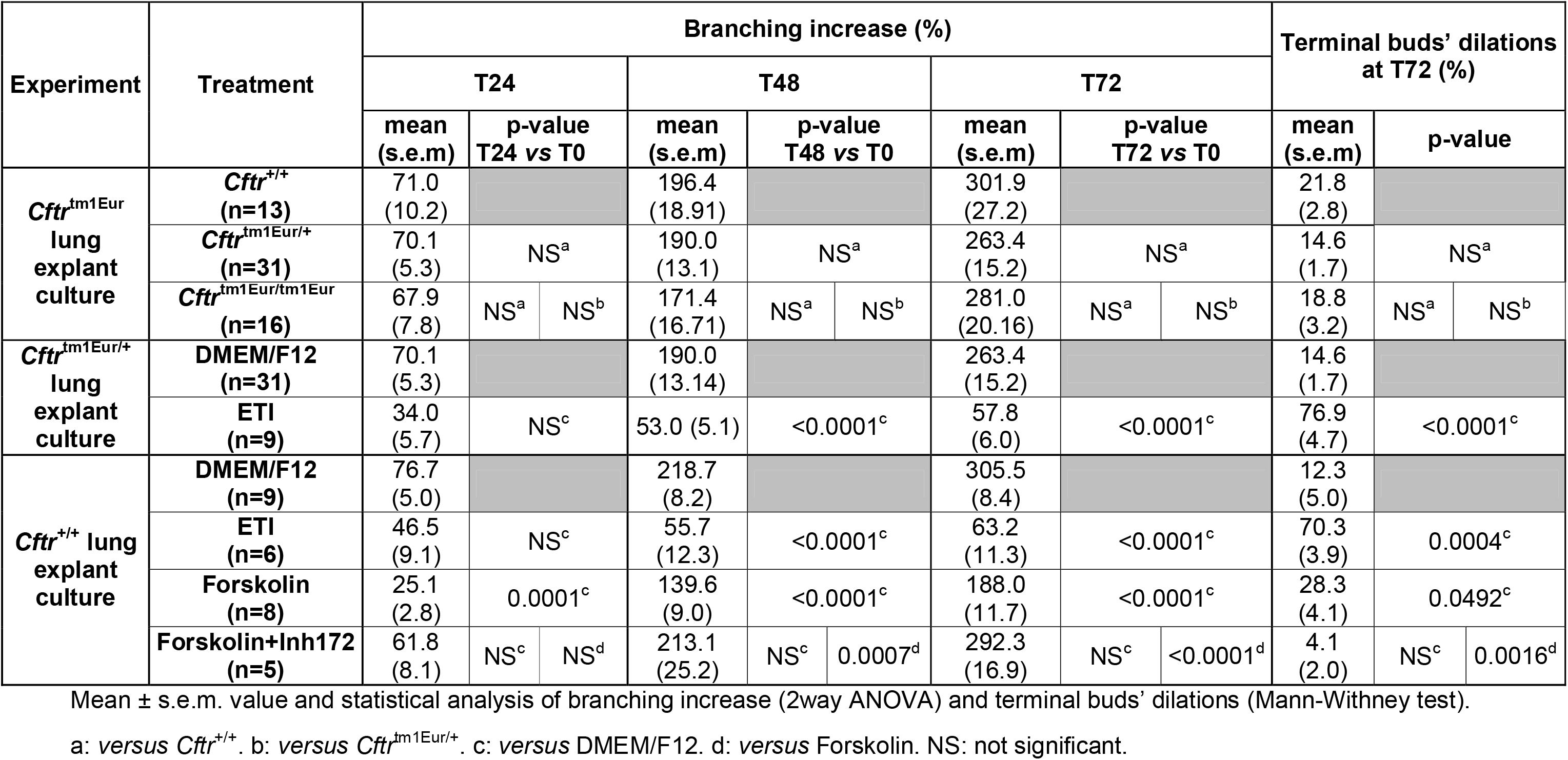
Lung branching and terminal buds’ dilations in WT *Cftr*^+/+^ and F508del heterozygous *Cftr*^tm1Eur/+^ lung explant culture.

### CFTR gain of function induced by ETI affects lung development in F508del heterozygous *Cftr*^tm1Eur/+^ mice

We investigated the effect of 3-day exposure to ETI on branching morphogenesis in lung explants from *Cftr*^tm1Eur/+^ embryos collected at E12.5. We first validated that *per se* DMSO, used as vehicle for ETI, has no effect on lung branching in culture when compared to media **(Figure S2**, **Table S2)**.

ETI incubation did not induce cytotoxicity as shown by the normal aspect of all lung explants and the easy distinction of the epithelial and mesenchymal layers. As shown in **Figure 2** and **Table 1**, there was a significant decrease in lung branching after 48h and 72h of culture for the *Cftr*^tm1Eur/+^ lung explants treated with ETI as compared to the control group incubated with media. The mean lung branching of the ETI group was decreased by 51% ± 8% at T24 (NS), 72% ± 3% at T48 (p<0.0001) and 78% ± 2% at T72 (p<0.0001). The percentage of terminal buds’ dilations of *Cftr*^tm1Eur/+^ lung explants was significantly increased by 5.3-fold (±0.3) after 72 hours (p<0.001).

**Figure 2.**
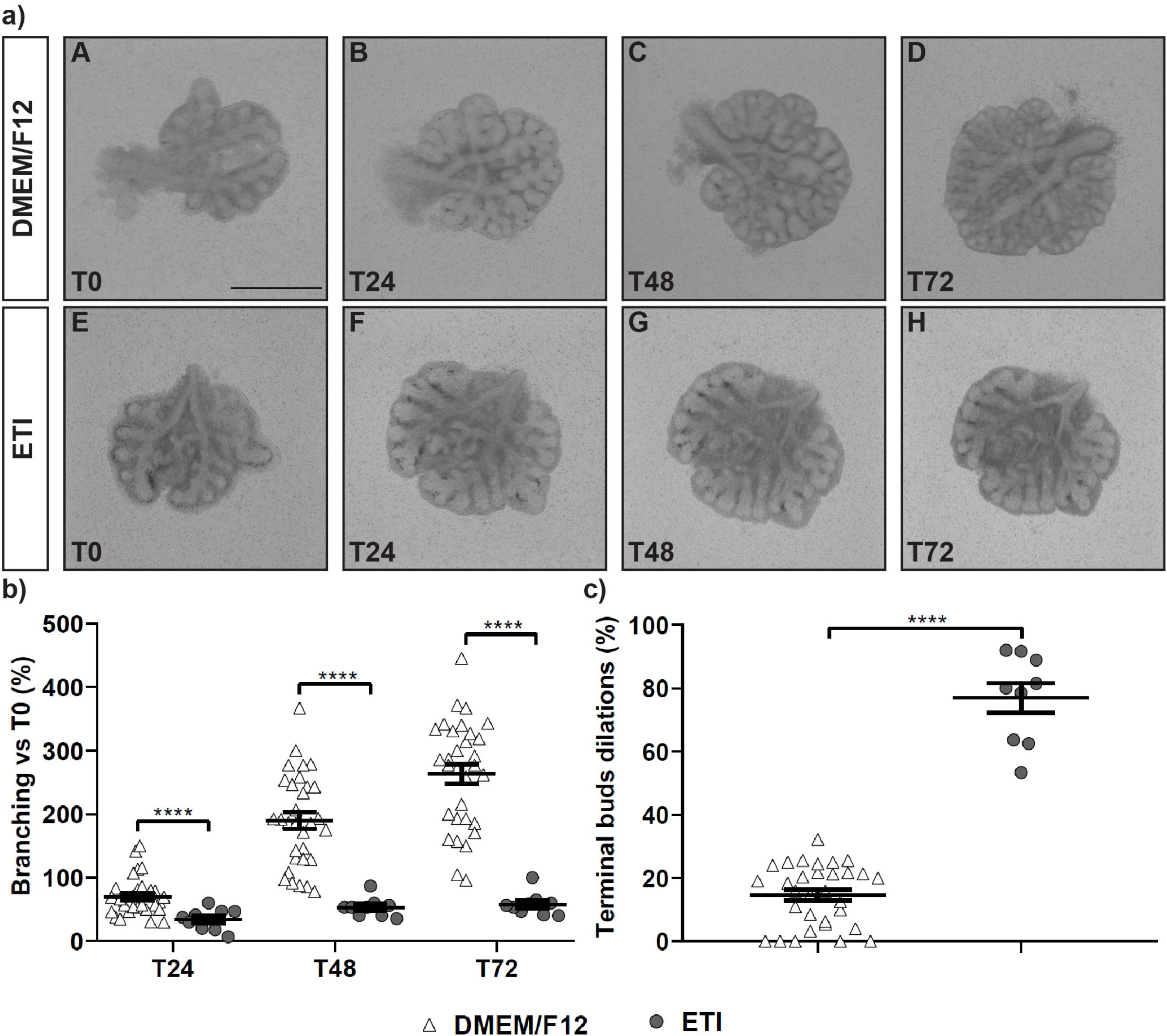
*Ex vivo* lung explants from E12.5 *Cftr*^tm1Eur/+^ embryos cultured with Elexacaftor/Tezacaftor/Ivacaftor association. A. Lung explants from *Cftr*^tm1Eur/+^ embryos cultured for 3 days in DMEM/F12 alone (A-D)(n=31), or DMEM/F12+ETI (E-H)(n=9). Scale bar: 1mm. B. Number of terminal buds at T24, T48 and T72, expressed as the percentage in branching increase compared to baseline (T0). C. Number of terminal buds’ dilations at T72, expressed as the percentage of the dilations increase compared to mean number of terminal buds’ dilations in the media group. Data are expressed as mean ± s.e.m. ****p<0.0001.

Altogether, these results suggest that acute ETI exposure of the lung at the pseudo-glandular stage negatively impacts on lung branching, and results in the formation of abnormal terminal dilations.

### ETI affects lung branching morphogenesis development in WT mice

To investigate the mechanisms underlying this pathogenic effect, we then asked whether the effect of ETI on lung branching might be related to the F508del mutation of *Cftr*. To this aim, we studied the effect of ETI on WT mouse lung explants at T24, T48 and T72 (**Figure 3**, **Table 1**). Lung branching was decreased upon incubation with ETI by 61% ± 12% at T24 (NS), 25% ± 6% at T48 (p<0.0001) and 21% ± 4% at T72 (p<0.0001) in comparison to the media control group. Similarly, there was a significant increase in the percentage of terminal buds’ dilations by 5.7-fold (±0.3) after 72h of culture for the ETI group when compared to the media control group (p=0.004). These changes were not significantly different from those observed in the *Cftr*^tm1Eur/+^ explants treated with ETI (**Table 1**) suggesting that the ETI triple combination affects lung branching morphogenesis independently of the presence of a F508del *Cftr* mutation.

**Figure 3.**
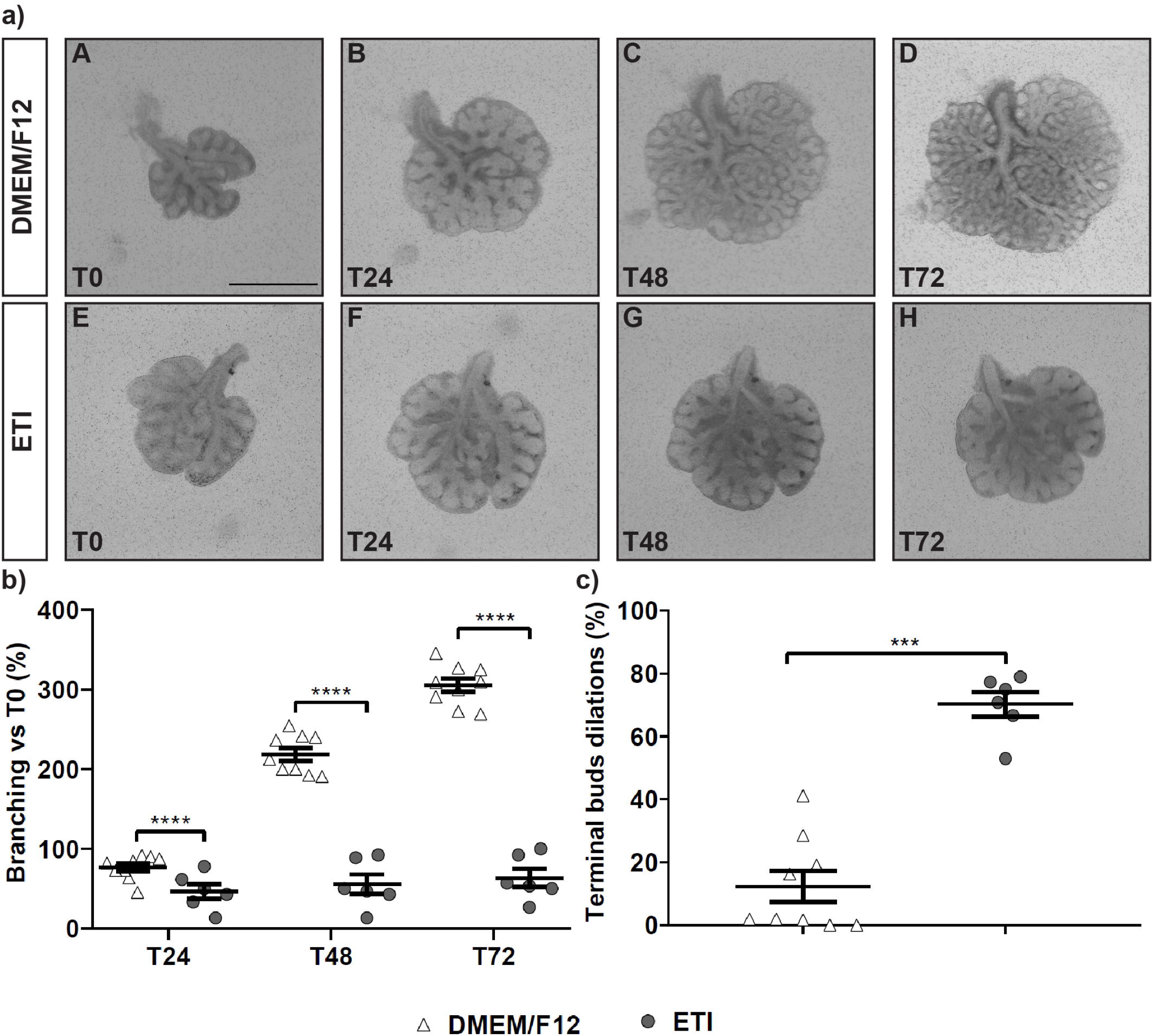
*Ex vivo* lung explants from E12.5 WT mouse embryos cultured with Elexacaftor/Tezacaftor/Ivacaftor association. A. Lung explants from WT embryos cultured for 3 days in DMEM/F12 alone (A-D)(n=9), or DMEM/F12+ETI (E-H)(n=6). Scale bar: 1mm. B. Number of terminal buds at T24, T48 and T72, expressed as the percentage in branching increase compared to baseline (T0). C. Number of terminal buds’ dilations at T72, expressed as the percentage of the dilations increase compared to mean number of terminal buds’ dilations in the media group. Data are expressed as mean ± s.e.m. ***p<0.001, ****p<0.0001.

### CFTR activation mediated by cAMP affects lung development in mice

As the mechanism of action of these drugs relies on the restoration of CFTR activity, we investigated whether the lung development defects observed after ETI exposure were associated to an increase in CFTR activity. To mimic this, we exposed lung embryos to the adenylyl cyclase activator Forskolin, which activates CFTR *via* PKA-dependent phosphorylation. We also checked the specificity of Forskolin action on CFTR by adding the CFTR specific inhibitor Inh172 in the culture medium (**Figure 4**, **Table 1**).

**Figure 4.**
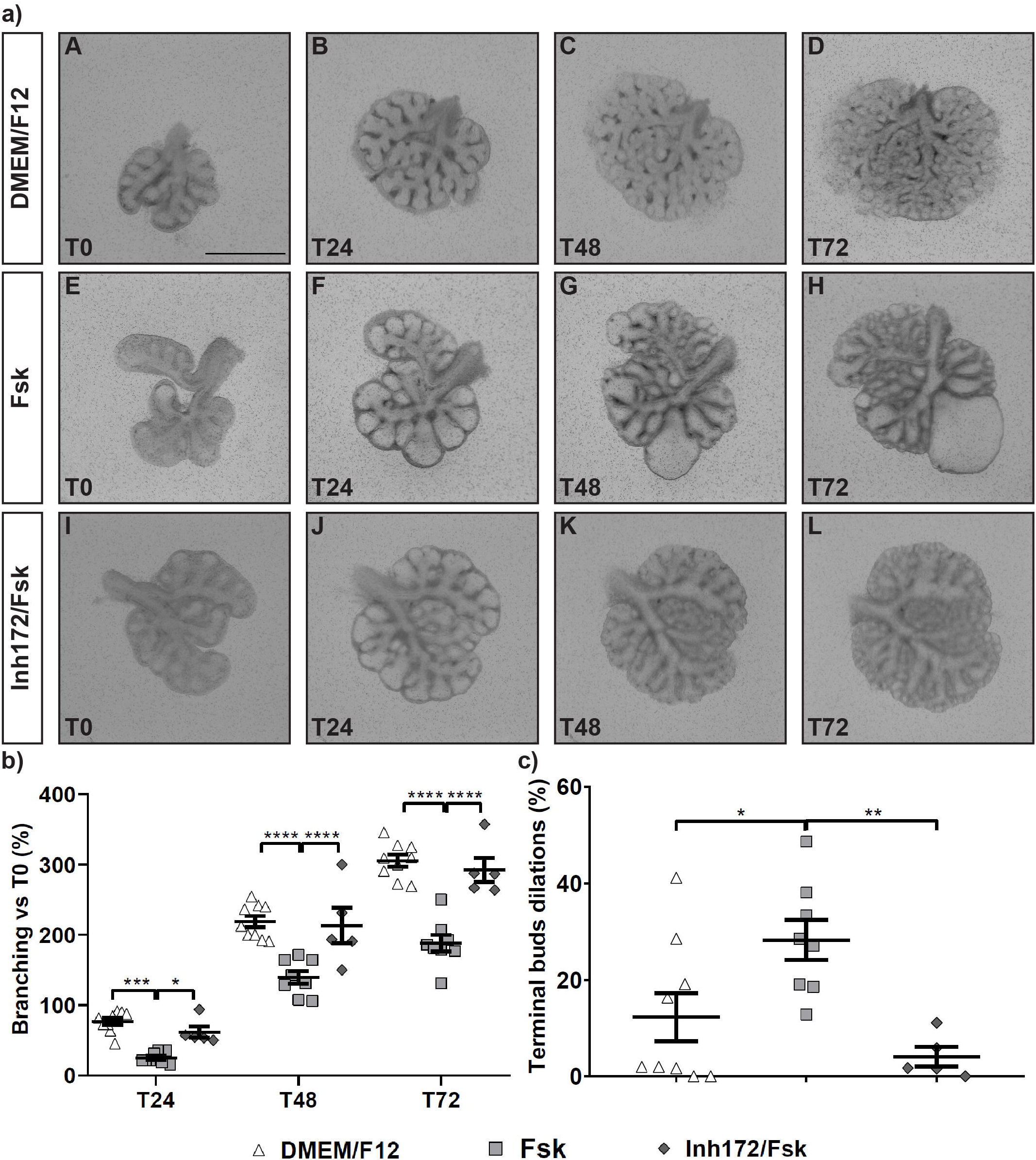
*Ex vivo* lung explants from E12.5 WT mouse embryos cultured with Forskolin or Forskolin+Inh172. A. Lung explants from WT embryos cultured for 3 days in DMEM/F12 alone (A-D)(n=9), DMEM/F12+Forskolin (E-H)(n=8), or DMEM/F12 +Forskolin+Inh172 (I-L)(n=5). Scale bar: 1mm. B. Number of terminal buds at T24, T48 and T72, expressed as the percentage in branching increase compared to baseline (T0). C. Number of terminal buds’ dilations at T72, expressed as the percentage of the dilations increase compared to mean number of terminal buds’ dilations in the media group. Data are expressed as mean ± s.e.m. *p<0.05, **p<0.01, ***p<0.001, ****p<0.0001.

We verified that *per se* EtOH, the vehicle of Forskolin, had no effect on lung branching compared to media (**Figure S2**, **Table S2**). We observed a significant decrease in lung branching after incubation with Forskolin of the lung explants at all time points tested. The mean lung branching was decreased by 67% ± 4% at T24 (p=0.0001), 36% ± 4% at T48 (p<0.0001) and 38% ± 4% at T72 (p<0.0001) in the Forskolin-treated group compared to the media control group. There was also a significant increase of the mean percentage of terminal buds’ dilations by 2.3-fold (±0.3) in the Forskolin-treated group (p<0.05).

Addition of Inh172 in combination with Forskolin to the culture medium rescued the lung branching phenotype and the mean of percentage of terminal buds’ dilations when compared to the media control group (**Figure 4**, **Table 1**). Altogether, these results show that the lung branching morphogenesis defects induced by ETI are recapitulated by the cAMP-dependent CFTR continuous activation.

### ETI modulates the expression of genes involved in foetal lung development in mice

We studied whether ETI modifies the expression of genes known to participate in fetal mouse lung development (**Figure 5**, **Table 2**). To do so, we assessed the expression of (i) Fibroblast Growth Factor 10 (FGF10), a chemoattractant crucial to lung branching and produced by clustered mesenchymal cells at sites where epithelial lung buds will elongate [23]; (ii) FGFR2IIIb, the FGF10 receptor on epithelial lung cells; (iii) Sonic Hedgehog (SHH) and Hedgehog-interacting protein (HHIP). SHH restricts FGF10 expression to the distal tips of the lung buds and, together with HHIP, inhibits FGF10 expression in the inter-bud regions, allowing localized new bud outgrowth [24].

**Figure 5.**
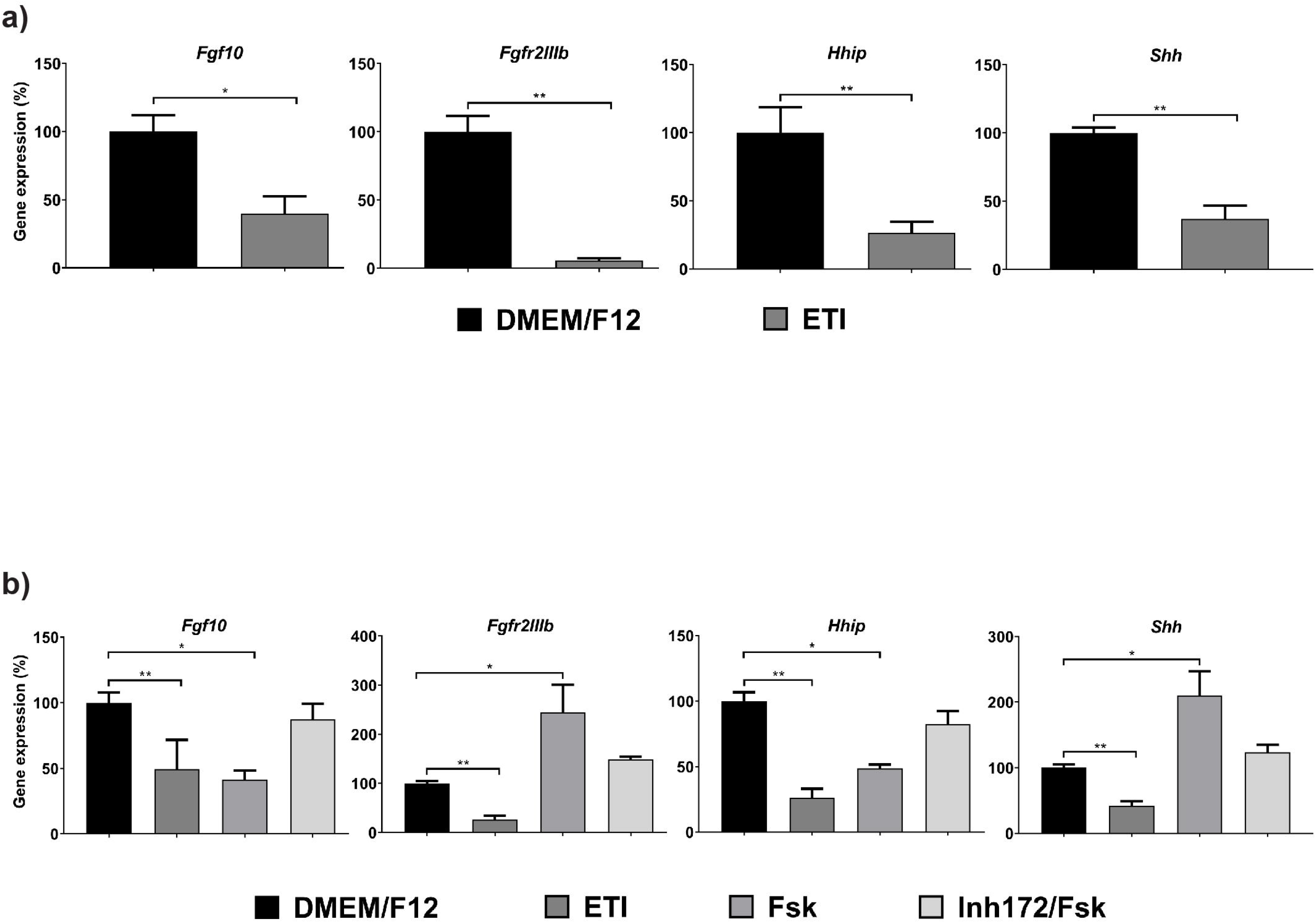
RT-qPCR gene expression of *Fgf10*, *Fgfr2IIIb*, *Shh* and *Hhip* in mice embryonic lung explant. A. Explants from *Cftr*^tm1Eur/+^ embryos cultured for 3 days in DMEM/F12 alone (n=5), or DMEM/F12+ETI (n=6). B. Explants from WT embryos cultured for 3 days in DMEM/F12 alone (n=5), DMEM/F12+ETI (n=5), DMEM/F12+Forskolin (n=3), or DMEM/F12 +Forskolin+Inh172 (n=4). Data are expressed as mean ± s.e.m. *p<0.05, **p<0.01.

**Table 2.**
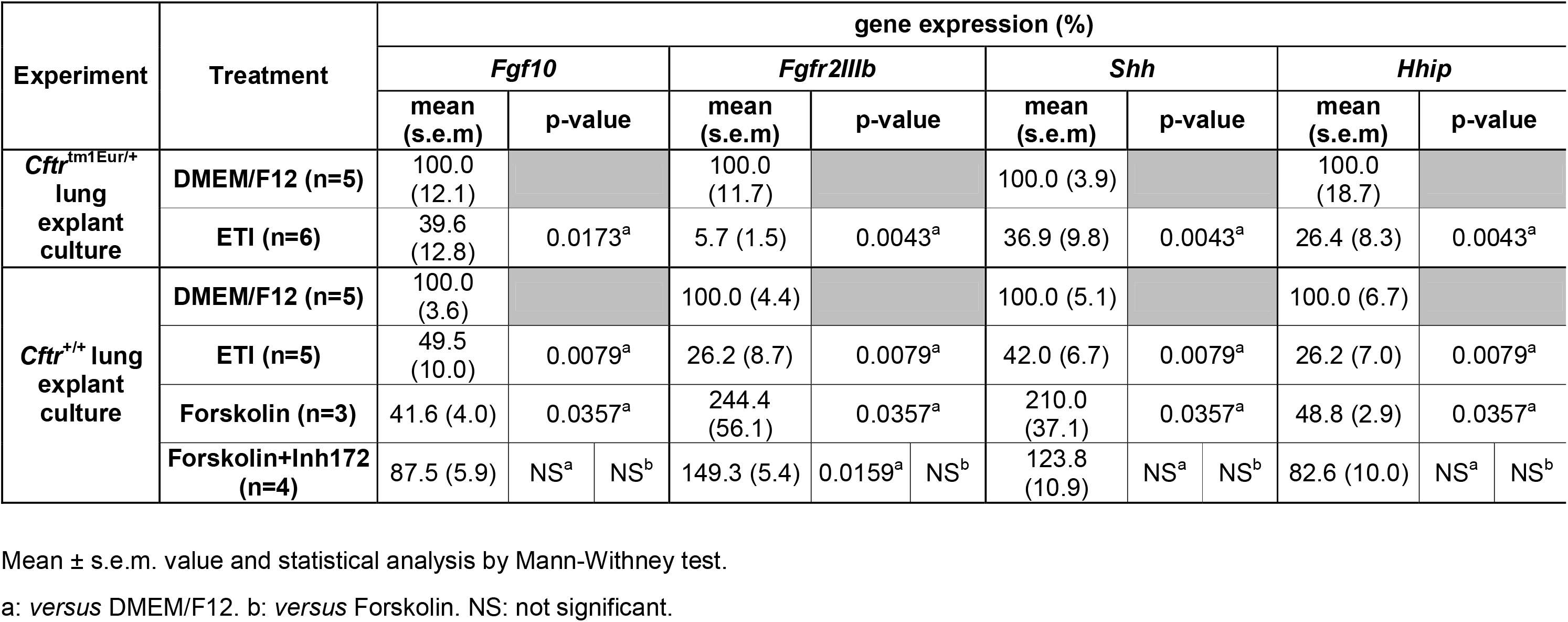
*Fgf10*, *Fgfr2IIIb*, *Shh* and *Hhip* expression in in WT *Cftr*^+/+^ and F508del heterozygous *Cftr*^tm1Eur/+^ lung explant.

RT-qPCR assays revealed that there was a significant decrease in *Fgf10*, *Fgfr2IIIb*, *Shh* and *Hhip* expression in lung explants from both E12.5 *Cftr*^tm1Eur/+^ and WT embryos treated by ETI when compared to the media-treated group (**Figure 5a**, **Table 2**).

The same expression profile was observed for *Fgf10* and *Hhip* in lung explants of WT embryos cultured in presence of Forskolin compared to the DMEM/F12 group (p=0.03) (**Figure 5b**, **Table 2**). This effect was partially reversed by the addition of Inh172. This contrasted with the increased expression of *Fgfr2IIIb* and *Shh* expression after incubation with Forskolin when compared to the media group (p=0.03 each). The expression of these two genes was partially restored to the control group level in the lung of WT embryos cultured with Forskolin+Inh172 but the difference was not statistically significant.

## Discussion

Our results provide evidence that the triple association of Elexacaftor/Tezacaftor/Ivacaftor alters lung morphogenesis of WT and heterozygous F508del mouse embryos at the pseudo-glandular stage by inducing lung branching defects and abnormal terminal bud’s dilation. This is related, at least partially, to CFTR activation as a similar morphological pattern is observed in lung from WT embryos cultured with the cAMP activator Forskolin, and this phenotype is partially reversed with the addition of the CFTR specific inhibitor, Inh172. These morphological changes are associated with decreased transcript levels of genes involved in mouse foetal lung development, including the growth factor *Fgf10* and its receptor *Fgfr2IIIb*, the ligand *Shh* and its effector *Hhip*. These observations made in a murine model suggest that ETI treatment may alter lung morphogenesis and induce the development of cystic lesions by inhibiting FGF10 signalling.

*CFTR* expression is spatially and temporally regulated in the human foetal lung [16, 25–30]. *CFTR* transcripts are detected strongly from the 12^th^ week of pregnancy at the pseudo-glandular stage, the period when the conducting airways and initial acinar framework develop, with a progressive increase up to the 24^th^ week followed by a very low expression after birth. Similarly to the human lung, we found in our mouse model, a gradual increase of *Cftr* transcript levels from the pseudo-glandular stage at E12.5, up to the canalicular stage at E18.5, further followed by a decrease from birth onwards. During lung development, CFTR is localized in the epithelium of the small airways [31], and is involved in foetal lung liquid secretion and lung organogenesis, possibly participating to the mechanico-sensory process regulating Wnt/β-catenin signalling [29]. Its specific inhibition in human foetal lung explants significantly decreases chloride and fluid active secretion [17]. However, defective expression of CFTR is not associated with abnormal lung organogenesis as the CF babies do not have abnormal lung airways and we did not observe a significant modification in lung branching and terminal bud dilations between embryos carrying the WT (*Cftr*^+/+^), the heterozygous *Cftr*^tm1Eur/+^ and the *Cftr*^tm1Eur/tm1Eur^ genotypes.

In this study, we show that the pharmacological activation of CFTR at the pseudo-glandular stage with ETI induces lung morphogenesis alterations characterized by major defects in lung branching and terminal bud dilation. We provide evidence that these defects are due to CFTR activation because similar defects are observed in WT embryonic lung explants treated with Forskolin, a phenotype reversed by the addition of the CFTR inhibitor Inh172. These results recall those obtained in the kidney epithelium where Forskolin increases the size and the number of cysts formed by Madin-Darby canine kidney cells expressing WT human CFTR, a pattern which was totally reversed by addition of CFTR Inh172 [32, 33]. This suggests that pharmacological activation of WT CFTR may contribute to the formation of epithelial cystic structures by increasing fluid secretion. In the context of F508del heterozygosity, this would combine an excessive and continuous activation of both the WT CFTR and of the corrected F508del CFTR.

Importantly, we observed that ETI consistently decreases transcript levels of both *Fgf10* and its receptor in epithelial cells *Fgfr2IIIb*. In the human lung, defective FGF10 and FGFR2IIIb expression are associated with defective lung branching and cyst-like terminal buds’ dilations [34]. Decrease in *Fgf10* expression was also detected after Forskolin exposure, and was partially rescued by Inh172, showing that this modification was related, directly or indirectly, to CFTR activation. In contrast, *Fgfr2IIIb* and *Shh* were decreased upon ETI exposure but were increased after Forskolin exposure. SHH signalling pathway is known to occur as a negative feedback mechanism during lung branching morphogenesis in mouse by inhibiting the local FGF10 signal [35]. The combined decreases in *Fgf10* expression and of its receptor after ETI exposure should ultimately lead to the total arrest of lung branching, whereas for Forskolin, increase in *Shh* might inhibit FGF10 signal competing with the remaining *Fgfr2IIIb* expression, which would maintain a partial lung branching. The decreased *Shh* expression observed in the lung explants of WT and heterozygous F508del *Cftr* embryos treated with ETI suggests that the arrest of lung branching seen with these lung explants could implicate additional molecular mechanisms that are still unknown.

The molecular mechanisms of action of the ETI drugs remain unknown. Our results suggest an interaction between CFTR activation mediated by ETI modulators and the FGF10-FGFR2IIIb signalling pathway. Further investigations are necessary to elucidate more precisely the molecular mechanisms involved in these developmental defects, keeping in mind that modulation of the FGF10 pathway may not only affect lung but also other organ morphogenesis, such as kidney.

There are many pregnancies expected to occur upon ETI treatment. Indeed, the improving effect of this therapy is so strong that more and more CF young women will undertake pregnancies. However, so far there are very few data on the human foetal and neonatal outcomes for pregnant women treated by Elexacaftor, Tezacaftor or Ivacaftor. In this context, although there are interspecies differences in the structural and functional features of the placenta, animal studies are helpful in predicting drug toxicity during pregnancy [15]. No teratogenicity was observed when pregnant rats and mice were challenged with at least 3 times the maximum recommended human dose of each drug independently although drug transfer across the placenta was observed [13]._However, even though those studies are able to detect obvious macroscopic agenesis or aplasia, they do not evaluate more subtle organogenesis defect such as formation of the bronchial tree.

In human, there are few pharmacological data about CFTR modulators during pregnancy. Trimble *et al* showed a significant higher concentration of Lumacaftor in the cord blood compared to the maternal plasma concentrations and an equivalent concentration of Ivacaftor [13]. Placental transfer of Ivacaftor was also observed in recent studies in CF animal models [19, 36, 37] and it is estimated to be around 40% ± 20%. The mean foetal lung concentration ratio was very high, around 200% of the plasma, much higher than into brain or cerebrospinal fluid (CSF) [19].

Currently, it is recommended that CF pregnant women stop their medication during pregnancy and breastfeeding, based on limited human data. Our observation of lung branching arrest and cyst formation in ETI exposed explants call for caution. Development of other organs such as kidney might be also affected as FGF10 activates several intracellular signalling cascades, resulting in cell proliferation and differentiation [38]. This strengthens the current recommendations, or at least argues for a close monitoring of those pregnancies.

## Supporting information

supplementary manuscript

## Take home message

The triple association Elexacaftor/Tezacaftor/Ivacaftor alters lung branching morphogenesis of murine F508Del heterozygous embryos. Those results argue for a close monitoring of pregnancies in women treated with this combo-therapy.

## Acknowledgements

We thank the Animal core facility of Institut Necker Enfants Malades.

---

**Figure S1.**
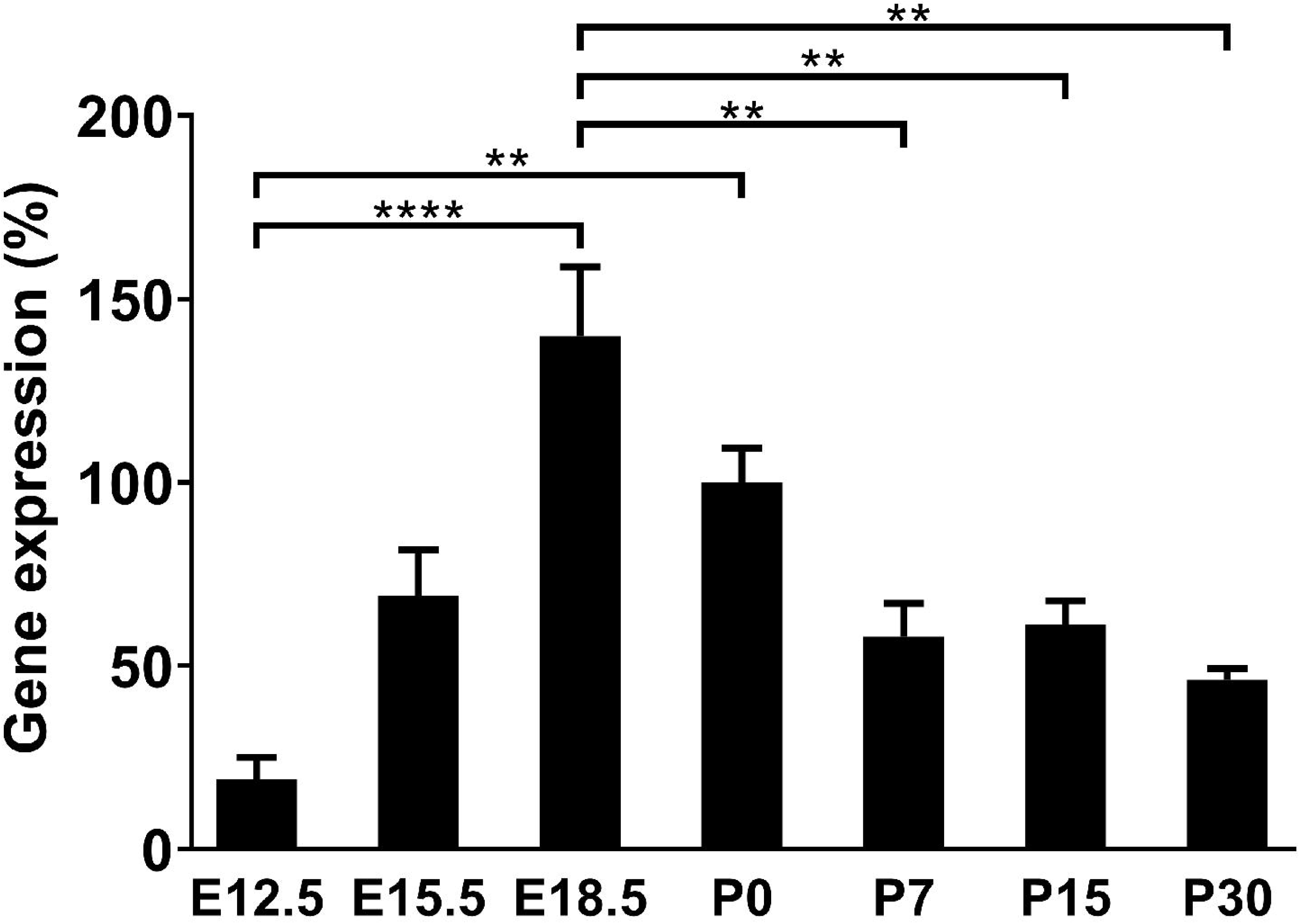
RT-qPCR *Cftr* expression during murine lung development. mRNA *Cftr* expression in lung from WT embryos (E12.5, E15.5 and E18.5) and newborns (P0, P7, P15 and P30). Data are expressed as mean ± s.e.m. **p<0.01, ****p<0,0001.

**Figure S2.**
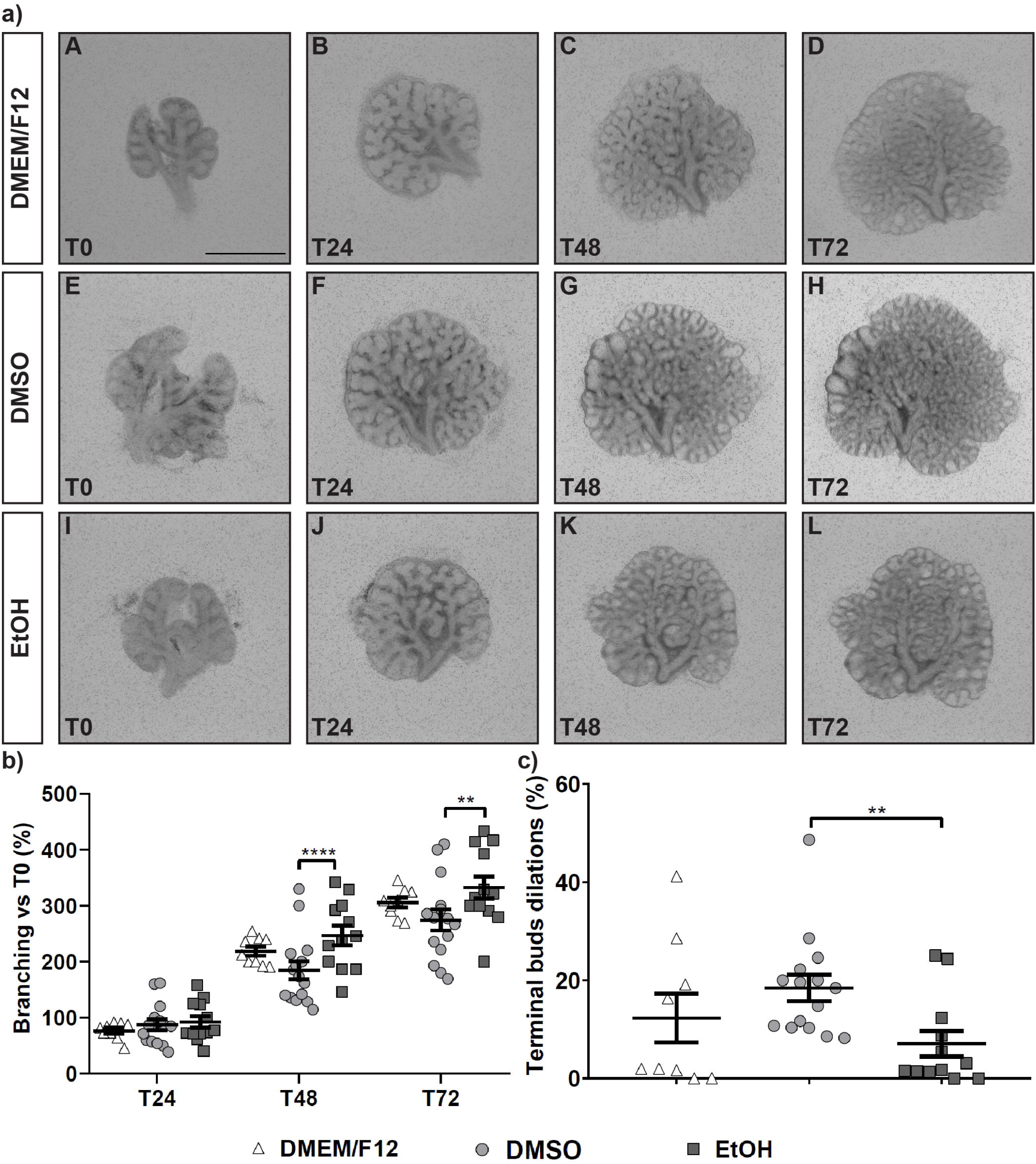
*Ex vivo* lung explants from E12.5 WT mouse embryos cultured with different drugs’ vehicle. A. Lung explants from WT embryos cultured for 3 days in DMEM/F12 alone (A-D)(n=9), DMEM/F12+DMSO (E-H)(n=15), or DMEM/F12+EtOH (I-L)(n=12). Scale bar: 1mm. B. Number of terminal buds at T24, T48 and T72, expressed as the percentage in branching increase compared to baseline (T0). C. Number of terminal buds’ dilations at T72, expressed as the percentage of the dilations increase compared to mean number of terminal buds’ dilations in the media group. Data are expressed as mean ± s.e.m. **p<0.01, ****p<0,0001.

**Figure S3.**
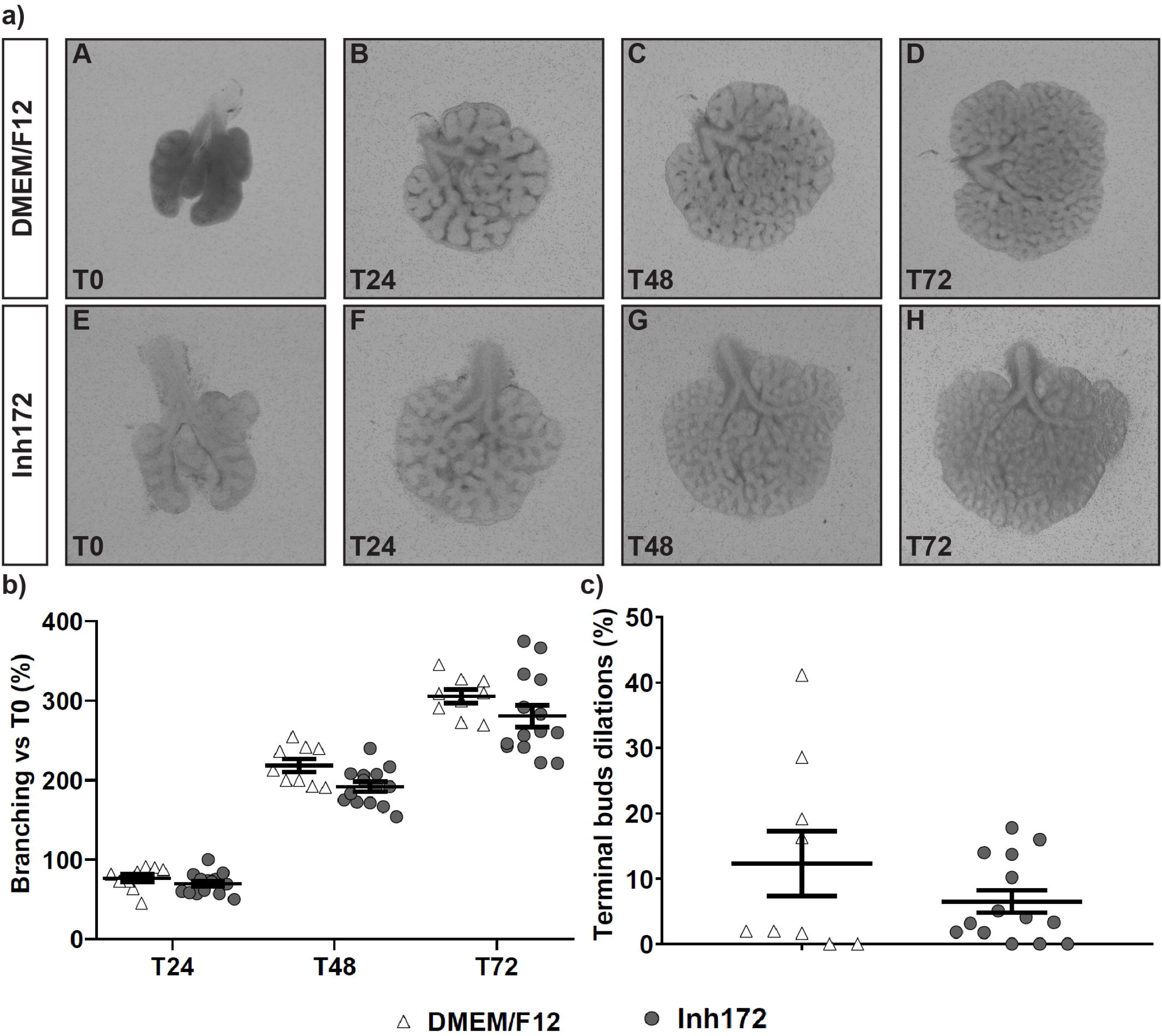
*Ex vivo* lung explants from E12.5 WT mouse embryos cultured with Inh172. A. Lung explants from WT embryos cultured for 3 days in DMEM/F12 alone (A-D)(n=9), or DMEM/F12+Inh172 (E-H)(n=14). Scale bar: 1mm. B. Number of terminal buds at T24, T48 and T72, expressed as the percentage in branching increase compared to baseline (T0). C. Number of terminal buds’ dilations at T72, expressed as the percentage of the dilations increase compared to mean number of terminal buds’ dilations in the media group. Data are expressed as mean ± s.e.m.

**Table S1:**
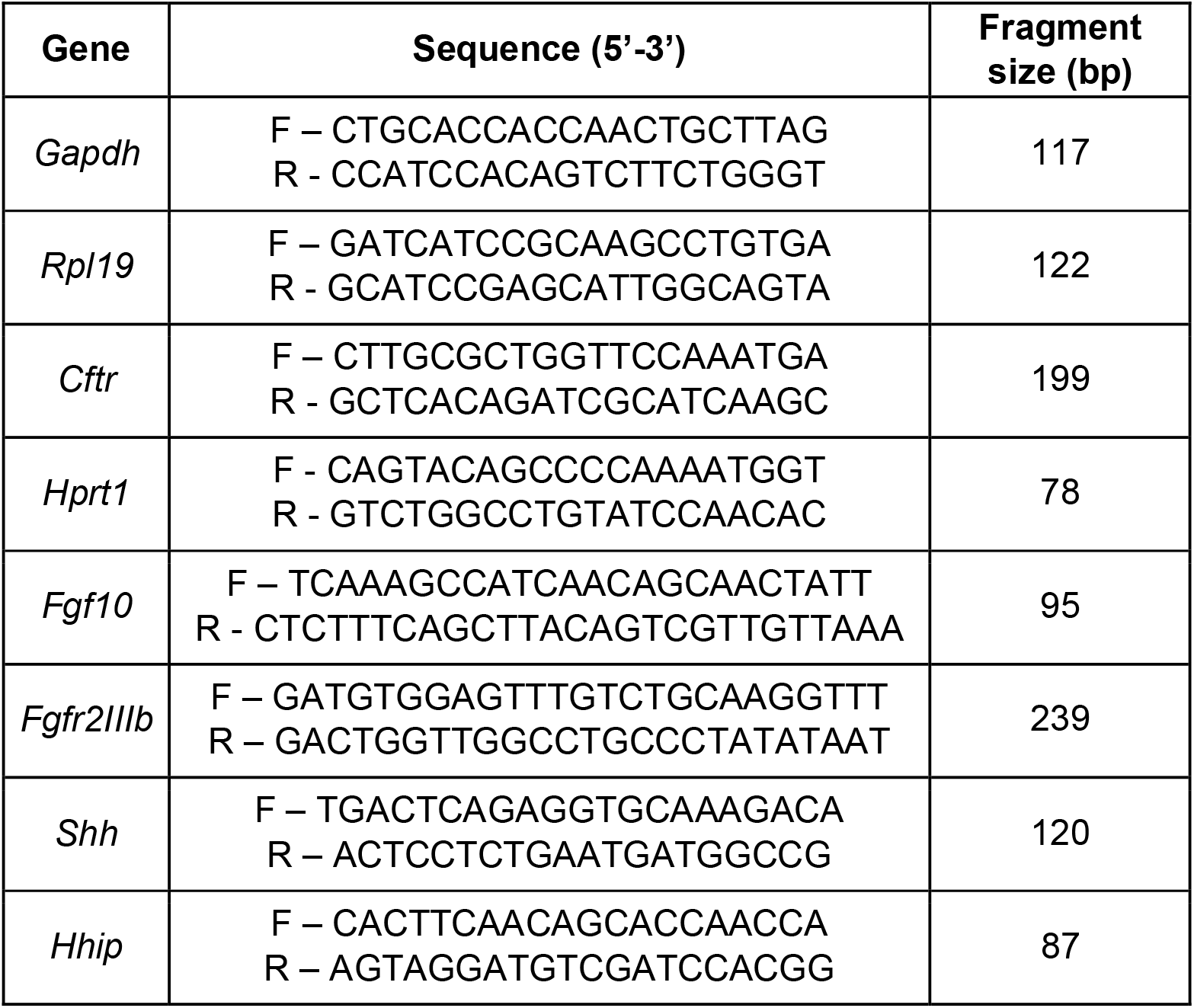
Primers sequences used RT-qPCR analysis.

**Table S2.** Mean value and statistical analysis of WT *Cftr*^+/+^ and F508del heterozygous *Cftr*^tm1Eur/+^ lung explant culture upon DMEM/F12 DMSO, EtOH, Inh172.

| Experiment | Treatment | Branching increase (%) |  |  |  |  |  |  |  |  | Terminal buds' dilations at T72 (%) |  |  |
| --- | --- | --- | --- | --- | --- | --- | --- | --- | --- | --- | --- | --- | --- |
|  |  | T24 vs T0 |  |  | T48 vs T0 |  |  | T72 vs T0 |  |  |  |  |  |
|  |  | mean (s.e.m) | p-value |  | mean (s.e.m) | p-value |  | mean (s.e.m) | p-value |  | mean (s.e.m) | p-value |  |
| *Cftr*<sup>+/+</sup> lung explant culture (control groups) | DMEM/F12 (n=9) | 76.7 (5.0) |  |  | 218.7 (8.2) |  |  | 305.5 (8.4) |  |  | 12.3 (5.0) |  |  |
|  | DMSO (n=15) | 87.3 (9.5) | NS<sup>a</sup> |  | 184.6 (15.6) | NS<sup>a</sup> |  | 274.5 (18.2) | NS<sup>a</sup> |  | 18.4 (2.7) | NS<sup>a</sup> |  |
|  | EtOH (n=12) | 92.5 (10.2) | NS<sup>a</sup> | NS<sup>b</sup> | 247.1 (17.7) | NS<sup>a</sup> | 0.0215<sup>b</sup> | 332.7 (19.9) | NS<sup>a</sup> | 0.0356<sup>b</sup> | 7.1 (2.6) | NS<sup>a</sup> | 0.0043<sup>b</sup> |
|  | Inh172 (n=14) | 69.5 (3.6) | NS<sup>a</sup> | NS<sup>b</sup> | 191.9 (6.2) | NS<sup>a</sup> | NS<sup>b</sup> | 280.6 (13.6) | NS<sup>a</sup> | NS<sup>b</sup> | 6.5 (1.7) | NS<sup>a</sup> | 0,0004<sup>b</sup> |

Mean ± s.e.m. value and statistical analysis of branching increase (2way ANOVA) and terminal buds' dilations (Mann-Withney test).
a: *versus* DMEM/F12. b: *versus* DMSO. NS: not significant.

