## supplementary manuscript for "Elexacaftor/Tezacaftor/Ivacaftor alters branching morphogenesis of the mouse embryonic lung"

**Real-time quantitative PCR analysis**

For RT-qPCR analysis, lung explants were snap-frozen in liquid nitrogen at T72. Lung total RNA was extracted with TRIzol™ (LS15596018, Invitrogen, Villebon-Sur-Yvette, France) according to the manufacturer’s instructions. cDNA was synthetized using High-capacity cDNA Reverse Transcription Kit (4368814, Invitrogen, Villebon-Sur-Yvette, France) with 1µg of total RNA and random primers. RT-qPCR was performed using the SYBR Green GoTaq qPCR Master Mix (A6001, Promega, Charbonnières-les-Bains, France) and a thermal cycler (qTOWER^3^, Biometra, Göttingen, Germany). Real-time quantification was achieved by measuring the increase in fluorescence caused by SYBR Green dye binding to double-stranded DNA at the end of each amplification cycle.

For *Cftr* expression during lung development, the expression was normalized using the geometrical mean from references *Rpl19* and *Gapdh* calculated with Pfaffl equation [1, 2].

The relative expression of *Fgf10*, *Fgfr2IIIb*, *Shh*, and *Hhip* were determined by using the ΔΔCt (threshold cycle) method of normalized sample (ΔCt) in relation to the expression of a calibrator sample used to normalize data, according to the manufacturer’s protocol. *Hprt1* mRNA was used as reference. Each PCR assay included a no-template control and a sample without reverse transcriptase. All measurements were performed in triplicates. Primers sequences are presented in Supplementary **Table S1**.

1.	Pfaffl MW. A new mathematical model for relative quantification in real-time RT-PCR. *Nucleic Acids Res.* 2001; 29: 45e–445.

2.	Vandesompele J, Preter KD, Roy NV, *et al*. Accurate normalization of real-time quantitative RT-PCR data by geometric averaging of multiple internal control genes. : 12.
